# Design, development, and preliminary assessment of a novel peripheral intravenous catheter aimed at reducing early failure rates

**DOI:** 10.1101/2022.06.20.496233

**Authors:** Barry J. Doyle, Lachlan J. Kelsey, Caroline Shelverton, Gabriella Abbate, Carmen Ainola, Noriko Sato, Samantha Livingstone, Mahe Bouquet, Margaret R Passmore, Emily S. Wilson, Sebastiano Colombo, Kei Sato, Keibun Liu, Silver Heinsar, Karin Wildi, Peter J. Carr, Jacky Suen, John Fraser, Gianluigi Li Bassi, Samantha Keogh

## Abstract

**Background:** Peripheral intravenous catheters (PIVCs) are the most commonly used invasive medical device, yet despite best efforts by end-users, PIVCs experience unacceptably high early failure rates. We aimed to design a new PIVC that reduces the early failure rate of in-dwelling PIVCs and we conducted preliminary tests to assess its efficacy and safety in a large animal model of intravenous access.

**Methods:** We used computer-aided design and simulation to create a PIVC with a ramped tip geometry, which directs the infused fluid away from the vein wall; we called the design the FloRamp™. We created FloRamp prototypes (test device) and tested them against a market-leading device (BD Insyte™; control device) in a highly-controlled setting with five insertion sites per device in four pigs. We measured resistance to infusion and visual infusion phlebitis (VIP) every six hours and terminated the experiment at 48 hours. Veins were harvested for histology and seven pathological markers were assessed.

**Results:** Computer simulations showed that the optimum FloRamp tip reduced maximum endothelial shear stress by 60%, from 12.7Pa to 5.1Pa, compared to a typical PIVC tip, and improved the infusion dynamics of saline in the blood stream. In the animal study, we found that 2/5 of the control devices were occluded after 24 hours, whereas all test devices remained patent and functional. The FloRamp created less resistance to infusion (0.73±0.81 vs 0.47±0.50, p=0.06) and lower VIP scores (0.60±0.93 vs 0.31±0.70, p=0.09) that the control device, although neither findings were significantly different. Histopathology revealed that 5/7 of the assessed markers were lower in veins with the FloRamp.

**Conclusions:** As PIVCs are used in almost every hospitalized patient, there is an urgent need to reduce failure rates. Herein we report preliminary assessment of a novel PIVC design, which could be advantageous in clinical settings through decreased device occlusion.

## INTRODUCTION

Peripheral intravenous catheters (PIVCs) are used to access the vasculature for the delivery of fluids and often life-saving medications, with up to 90% of patients requiring one during their hospital stay (1). PIVCs are the most widely used invasive medical devices (2), with over 2 billion purchased globally each year (3) and the market set to exceed $8 Billion by 2028 (4).

Although they play a significant role in medical practice and are used extensively, up to 50% of PIVCs fail within days once they are inserted into the vein (1, 5): they occlude, or diverge into the neighbouring tissue and must therefore be withdrawn before fulfilling their clinical use.

To address this problem of PIVC failure, research and practice has focused on optimising insertion and maintenance care, often encompassed within ‘care bundles’ (6). In a recent large randomised clinical trial (7), a team implemented a multimodal approach that included: best-practice hygiene and aseptic measures; insertion, maintenance and removal of PIVCs, along with comprehensive education for health-care professional and patients. During the 12-month test period, they implemented this bundle in conjunction with a leading PIVC (B.Braun Introcan Safety® (non-winged) device) in a population of more than 2,000 patients. across seven hospitals. As a result of this dedicated approach, they were able to reduce failures by 9%; but importantly, 37% of PIVCs failed despite their best efforts.

Commercially, there is a plethora of differently designed PIVCs available to clinicians. However, other than the PIVC’s polymeric formulation (8, 9), the invasive segment of the catheter has not changed in the last few decades. In addition, manufacturers have sought to improve the device through the addition of wings and device stabilisers (10). However, to improve PIVC functionality, the mechanics of the cannula interacting with the vein and the blood flow cannot be overlooked, especially when considered alongside Virchow’s Triad of venous thrombosis (11, 12). In the Triad, there are three factors involved: (1) endothelial injury; (2) abnormal blood flow, turbulence or blood stasis; and (3) hypercoagulability. While hypercoagulability is patient-specific and cannot easily be incorporated into a design brief, endothelial injury (i.e. damage) and turbulence or stasis (i.e. flow) are factors that are directly influenced by the PIVC, and thus can be improved through innovative device design.

Endothelial cells line the inside of veins and respond rapidly and sensitively to the mechanical conditions caused by blood flow (13). Shear stress, the frictional force of blood acting on the endothelium, is a fundamental mechanical force resulting from the flow of blood. At physiological levels, shear stress helps control the presentation of tissue factor on the surface of vascular cells, thus ensuring a normal coagulative balance. Supra- or sub-physiological levels of shear stress yield a procoagulant state in the vasculature. Hence, excessively high, or low shear stress, can predispose patients to venous thrombosis. Exposure to high shear stress stimulates endothelial cell cilia disassembly and disrupts vascular stability (14), and we discovered that when the PIVC is not optimally positioned in the vein and is directing fluid towards the vein wall, shear stress can reach levels which are 3,000 times over the physiological norm (15). Even if the PIVC is perfectly positioned in the vein, we found the shear stress to reach levels at least 13 times higher than the normal physiological level of the vein (16).

Irregular or abnormal blood flow through turbulence or stasis promotes endothelial expression of adhesion molecules and monocyte attachment to the cells (17). When a PIVC is inserted into a vein, the venous blood must then flow around the device, often competing with fluids infusing from the PIVC, which creates abnormal flow patterns. When a PIVC infuses directly along the vein wall, it also creates large regions of recirculation (16), especially if the PIVC is positioned proximal to a venous valve. When the surrounding endothelial cells detect this abnormal blood flow, the vessel response is phlebitis, a critical step in a cascade of events that leads to thrombosis and occlusion. Furthermore, highly disturbed flow activates platelets, widely considered to play a key role in thrombosis (18). Activated platelets clump together and stick to injured surfaces, such as an injured vessel wall.

Thus, there is overwhelming evidence that PIVC design needs improvement to reduce the unacceptable failure rates observed clinically. Our hypothesis is that incorporating the controllable aspects of Virchow’s Triad into the PIVC design will lead to better device functionality and reduced failure rates. Therefore, the aim of this study was to design a new PIVC and demonstrate proof of concept in a large animal model of intravascular (IV) access.

## METHODS

### Virtual Prototyping

#### Computer-aided design

Using the computer-aided design (CAD) tools in STAR-CCM+ (v12, Siemens, Germany), we simulated the geometries of a typical 20 Gauge (G) PIVC with an externally tapered tip, using dimensions implemented in previous investigations (16); outer diameter 1.1 mm, inner diameter 0.8 mm. This acts as a typical geometry to which we compared novel PIVC designs. We constrained our design with the following criteria:

- The design must reduce the shear stress applied to the vein around the PIVC tip during infusion;
- The design must improve infusion dynamics and promote diffusion of fluid into the bloodstream;
- The design must be as simple as possible to facilitate large-scale production; and
- The design must allow similar clinical application to existing PIVCs.

Through an iterative approach, we designed a novel asymmetric ramped tip that would ramp the infused fluid towards the centre of the vein (and thus the venous blood flow) and spare the endothelium from excessive shear stress. We named this design the FloRamp™, the geometry of which is shown in Figure 1A. We then investigated the optimal ramp geometry by modifying the ramp dimension to minimise shear stress without adversely directing the infused fluid into the superior vein wall.

**Figure 1:**
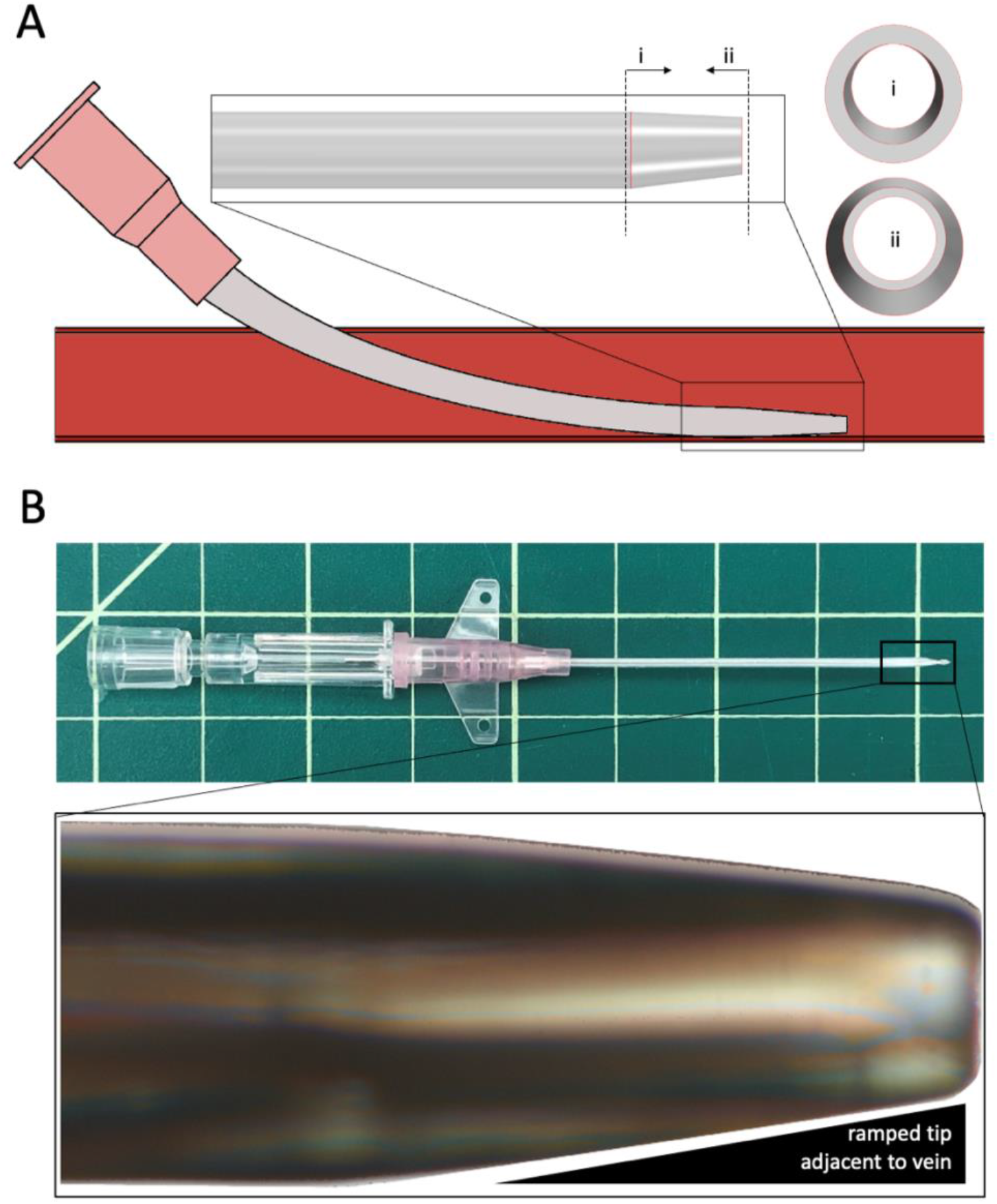
Prototyping of the FloRamp. (A) The computer-aided design showing the asymmetric ramped tip, with the insets showing views from within (i) and from outside (ii) the device. (B) Physical prototype produced.; in the background, each box is 12.5 × 12.5 mm. Insert shows asymmetric tip imaged using a light microscope.

#### Computer simulations of infusion

We applied methods detailed in our recent publication (16). Briefly, we computationally inserted the PIVCs into a 100 mm long vein of 3.3 mm diameter. The PIVC was inserted into the vein at an insertion angle of 20° before resting in the vein in an optimum configuration, with the majority of the device intravascular. We then infused a bolus of saline (NaCl 0.9%) at a rate of 1 ml/s into venous blood flow of 28 ml/min (velocity of 11 cm/s), with the blood set at 37°C and the saline set at 20°C. We ran all simulations on 512 cores of the Magnus supercomputer (Pawsey Supercomputing Centre, Perth) and extracted the resulting shear stress acting on a 2 cm section of vein immediately downstream of the PIVC tip after completion of each simulation. We also analysed the overall distribution of shear stress in the vein and the diffusion of saline into the vein during infusion.

### Physical Prototyping

After virtual prototyping, we produced the optimal FloRamp geometry. All physical prototyping was performed at a specialised catheter manufacturing facility (Medical Murray, IL, USA). Similar to a previous study (19), we decided to modify a commercial device to allow a true comparison of the FloRamp with an existing PIVC. We chose the 20 G BD Insyte™ winged 45 mm long catheter (Becton Dickenson, Franklin Lakes, NJ, USA), as the BD Insyte is widely used and familiar to most infusion therapists. Furthermore, by modifying an existing device, we can determine the true impact of the design, eliminating any differences due to material formulations, hub design, or other design factors that may contribute to failure.

We designed and fabricated a glass tipping dye with the required asymmetric offset. We then placed the BD Insyte cannula into the dye, ensuring the offset was positioned correctly which is confirmed by the positioning of the winged hub, before heating the dye under pressure, enabling the tip of the cannula to conform to the new asymmetric shape (Figure 1B). Because the FloRamp tip has a slightly smaller outlet diameter, we paired a 22 G needle (B.Braun, Germany) with the 20 G FloRamp. All prototypes were inspected for visual defects such as air bubbles, and after passing internal quality controls, they were packaged for sterilisation.

### Animal Study Design and Protocol

All procedures in this study were approved by the Queensland University of Technology (QUT) animal ethics committee (UAEC #2000000023) and all research involving animals followed the NIH Guide for the Care and Use of Laboratory Animals. We used four white female pigs (weight 35±2.3 Kg), all less than 1 year old. The animals were procured locally (Medina Pastoral Piggery, QLD, Australia) and were transferred to the QUT Medical Engineering Research Facility on the campus of The Prince Charles Hospital. The pigs were acclimatised in the animal house for seven days before the study. The experimental protocol was designed to mimic clinical practice. However, to eliminate potential movement of the device *in situ* and provide robust comparisons of the PIVC designs, all animals were sedated throughout the 48 hour study duration.

Prior to the study, the pigs fasted for 12 hours with free access to drinking water. On the first day of the study, pigs were anaesthetised and ventilated before being placed in the prone position in the operating room. The marginal veins of the ears of each pig (Figure 2A) were randomly assigned to either a FloRamp (test) or a BD Insyte (control), and the cephalic veins (Figure 2B) of a pig were also assigned to either device. The outer surface of the ears was clipped, and the ear skin was wiped with an antiseptic-soaked sponge and allowed to dry. Cephalic veins were accessed along the cranial surface of the radius on the right and left forelimb. A tourniquet was then applied at the base of the ear or around the forelimb at the level of the proximal radius to improve vein filling and increase venous dilation.

**Figure 2:**
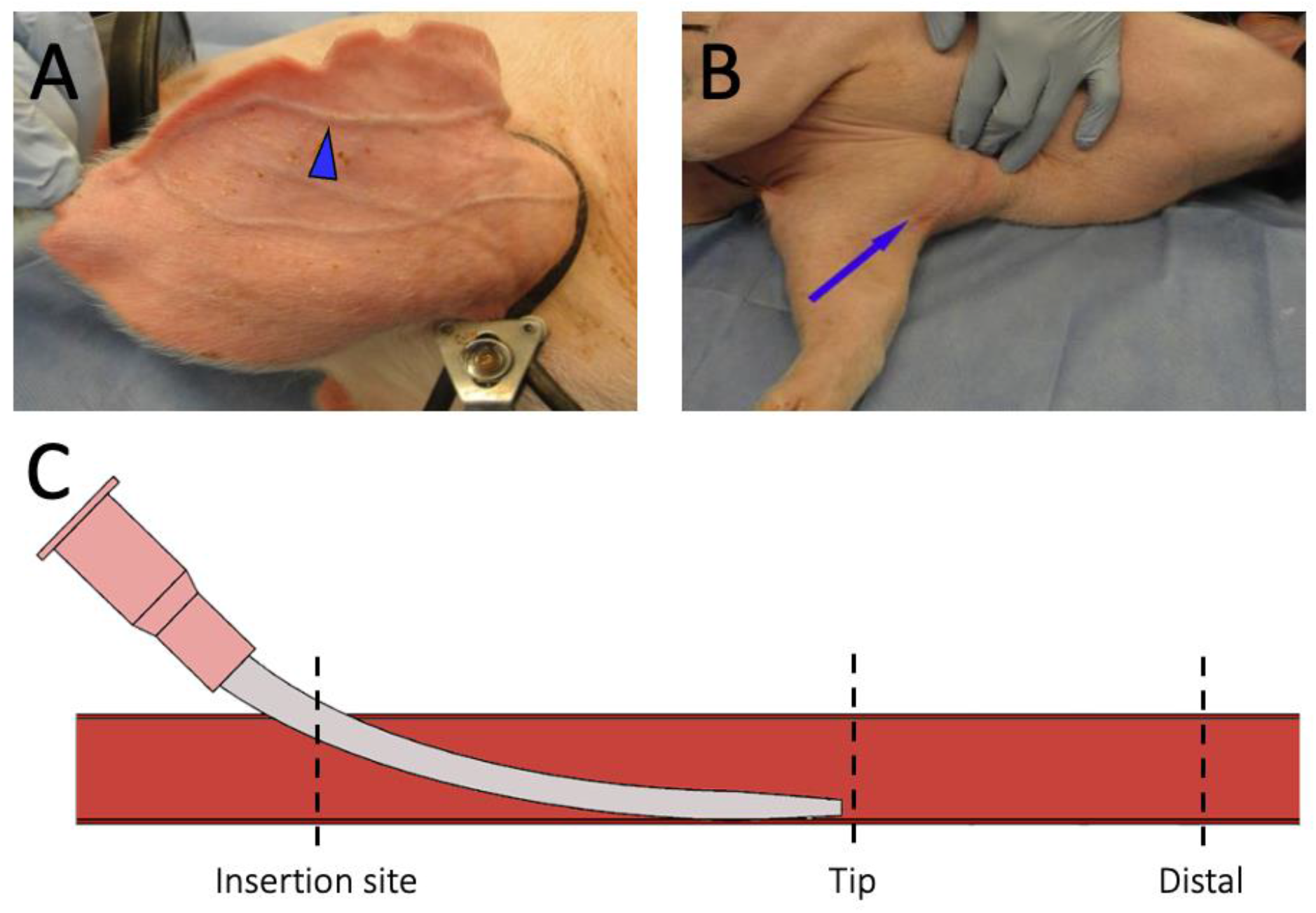
Pigs were sedated, ventilated, placed prone, and continuously monitored throughout the study, with measurements and saline flushes every six hours. (A) Marginal vein (arrow) of the ear and (B) cephalic vein used for vascular access. (C) Approximate locations at the insertion site, PIVC tip and distal to the PIVC, where samples were extracted for histopathology.

Because the FloRamp and BD Insyte are visually the same, clinical staff were unaware of which device they were inserting. PIVCs were inserted using standard techniques at an insertion angle of approximately 20°. Once a flashback was observed, the cannula was advanced to the centre of the vein, before the introducer needle was slowly removed. An IV link (MicroClave Neutral Connector, ICU Medical, CA, USA) was connected to the hub of the device and the device was flushed with heparinised saline. The hub of each device was secured to the skin with tissue adhesive (Histoacryl, B.Braun, Germany), the tourniquet was removed, and a sterile transparent dressing (Tegaderm, 3M, MN, USA) was placed over the insertion site. A custom-made ear brace was then placed into the inside of each ear and secured with tape (Nexcare, 3M, MN, USA).

A continuous infusion (to keep vein open, TKVO) of saline at 0.5 ml/Kg/hour was administered throughout the study. Every six hours, research nurses with more than 5 years of clinical experience recorded the Visual Infusion Phlebitis (VIP) score (20) following accepted clinical practice to assess phlebitis and potential occlusion. TKVO was then stopped and 10 ml of NaCl 0.9% was flushed from the central infusion line at 1 ml/s before the TKVO infusion was resumed. We previously deemed a flush rate of 1 ml/s to be a conservative rate applicable in clinical practice (15). Using clinically accepted grading, the VIP score ranged from 0 to 5 in ascending order of severity. In addition, staff graded resistance to infusion, where 0 meant no resistance to infusion, 1 meant some resistance to infusion, and 2 meant occlusion. Animals were monitored continuously over the entire study.

At the end of the study, pigs were euthanised with IV injection of 0.5ml//Kg Lethabarb via central cannula while under general anaesthesia. For histopathology samples, the ears and the cephalic veins of each pig were harvested via microsurgery and were fixed in 10% buffered formalin for at least 48 hours, processed and embedded in paraffin. Sections (5 µm) obtained from the PIVC tip region, the insertion region and distally from the tip (Figure 2C), stained with haematoxylin and eosin and examined by light microscopy by an independent trained pathologist (QML Vetnostics, QLD, Australia) blinded to the slide identity. We followed the methods described in Weiss et al. (19) and assessed samples for mural inflammation, thrombus formation, exfoliation of the endothelium, intimal proliferation, intimal oedema, intimal necrosis and mural haemorrhage. Each marker was graded on a scale of 0 to 4 (0 = none; 1 = slight; 2 = mild; 3 = moderate; 4 = severe) (21).

### Statistical Analysis

All measured data were collected on a study report form, collated and analysed in Microsoft® Excel (v16). Descriptive statistics are used to summarise subject and device characteristics, and outcomes are compared using Student’s t-test, with p<0.05 deemed significant.

## RESULTS

### Effects on infusion dynamics and endothelial shear stress

The optimum ramp geometry was a trade-off between reducing endothelial shear stress and improving infusion dynamics; in particular, we did not want to direct the infused fluid into the upper vein wall. We found the ideal geometry to be a ramp size of 0.125 mm off the horizontal plane (Figure 3A), whereas 0.15 mm increased the velocity of the infusion and created an undesirable recirculation region downstream of the tip, below the saline fluid jet. Therefore, we deemed a ramp size of 0.125 mm to yield an adequate reduction in shear stress, while ensuring infused fluids remain central to the venous flow (Figure 3B). This ramp size reduced the maximum endothelial shear stress by 60%, from 12.7 Pa to 5.1 Pa, compared to a PIVC with constant internal diameter (Figure 3C).

**Figure 3:**
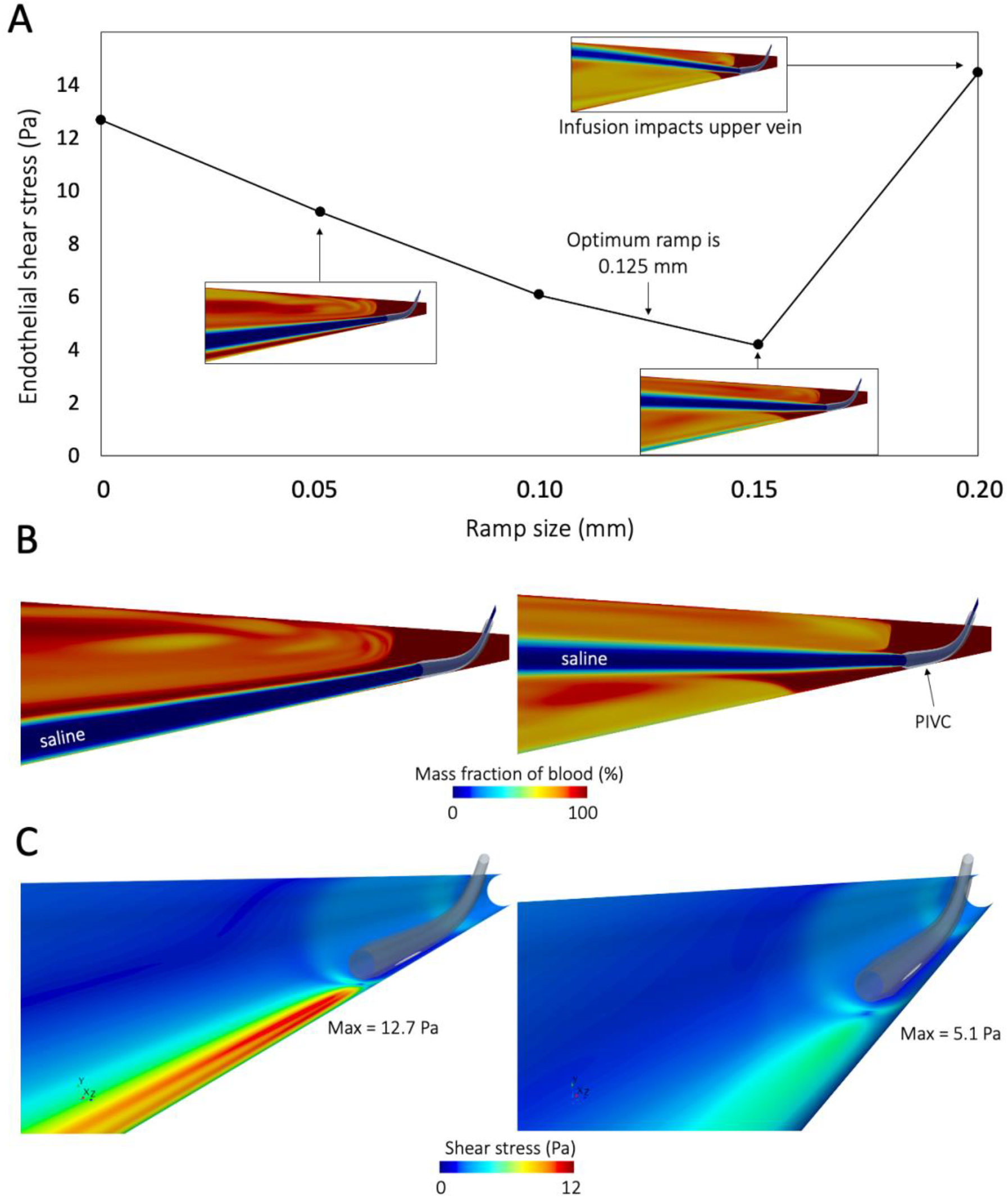
(A) Investigation of the ramped tip geometry determined that a ramp size of 0.125 mm is ideal as it reduces shear stress while ensuring optimum infusion dynamics, without impinging on the upper vein wall. (B) Diffusion of saline into the blood stream using the optimum tip geometry. Image shows the mass fraction of blood in the computer simulation during the bolus flush. (C) The optimum ramp geometry reduces the maximum shear stress on the endothelium by 60% compared to a standard PIVC tip.

### Animal Study

All four animals completed the 48-hour study and performance of tested cannulas was followed throughout the period of investigation.

### Resistance, VIP and failure rates

A key observation from the study was that 40% (2/5) of the control PIVCs became fully occluded during the 48 hour time period (Figure 4A). Despite using a continuous background infusion (i.e. TKVO), these two failed devices became occluded between 18-24 hours after insertion and the start of infusion therapy (i.e. TKVO). Although some resistance to infusion was observed, all FloRamp devices remained fully functional throughout the 48-hour study.

**Figure 4:**
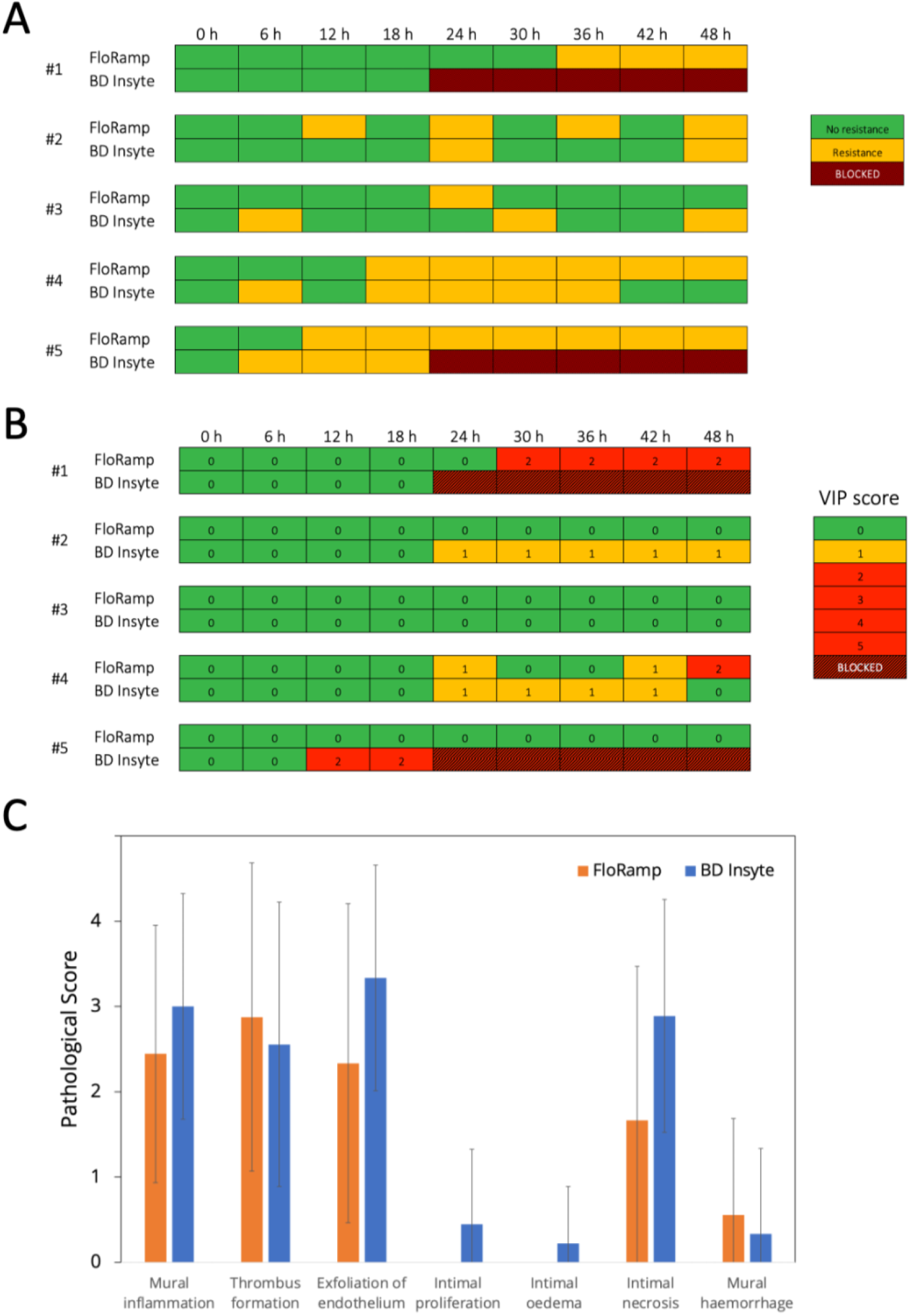
Observations from the animal trial of five comparative experiments of test devices (FloRamp) versus control devices (BD Insyte). Experiment #3 was a cephalic vein with the others being marginal ear veins. (A) Measures of resistance taken every 6 hours show that resistance to infusion generally began earlier in control PIVCs and that 40% of control devices (2/5; Experiments #1 and #5) became completely occluded after 24 hours *in vivo*. (B) VIP score represents a visual marker of inflammation (thrombophlebitis) and the test devices had reduced VIP scores compared to the controls. (C) Mean ± SD histopathology markers showing 5/7 markers were lower for veins with test PIVCs. Only 3/5 veins were used for comparative histology as we excluded comparisons with the two veins that occluded.

A trend toward lower mean resistance was found in the test group compared to the controls (0.73±0.81 vs 0.47±0.50, p=0.06). As shown in Figure 4A, in some veins both the test and control PIVCs experienced intermittent resistance throughout the 48-hour duration. Resistance was typically observed earlier in control devices; at 6 hours for 60% (3/5) of devices. Whereas the earliest sign of resistance in a test device was 12 hours (2/5 devices).

Mean VIP was lower in veins inserted with the test PIVCs compared to the controls, but such difference was not found to be statistically different (0.60±0.93 vs 0.31±0.70, p=0.099) (Figure 4B), with visual markers of inflammation observed earlier in control devices. Sixty percent (3/5) of veins inserted with the test PIVC did not experience any VIP score compared to 20% (1/5) of veins inserted with a control PIVC.

### Histopathological findings in cannulated veins

Pathological scores were lower in 5/7 of the markers assessed in veins inserted with a test device (Figure 4C), with differences in mural inflammation, exfoliation of endothelium and intimal necrosis of 19%, 30% and 42%, respectively. Veins inserted with test devices did not initiate any intimal oedema or intimal necrosis. Thrombus formation was marginally higher in test device veins as was mural haemorrhage, however mural haemorrhage was only observed at one tip location in one control device vein (1/9 sites), and two tip locations in one test device vein (2/9 sites).

## DISCUSSION

Despite best efforts by the end-user, the premature failure of IV cannulas remains at almost 40%, which is unacceptably high (7). Aligning with strong calls from the community for a change to the device itself (22), we set out to design an IV cannula that reduces the early failure of in-dwelling PIVCs and through a pilot study, explored efficacy and safety in a highly-controlled large animal model of IV access, using a protocol that closely mimics clinical practice. The key finding of our study is that a novel design modification to the PIVC tip reduces the maximum endothelial shear stress acting on the vein by 60%. By directing the infused fluid into the centre of the venous flow, the saline diffuses further downstream and away from the endothelium, the sensitive endothelial cells are spared. This reduction in injury and improvement in infusion dynamics potentially translates to an advantage over current PIVCs through a reduction in occlusion rate.

The most important clinical implication of our study is the reduced failure rate of the FloRamp. We compared the FloRamp to one of the most widely used PIVC on the market that is typical in design and material to most used in clinical settings. We observed two of the five control PIVCs to become fully occluded within 24 hours of insertion. Despite the obvious small sample size, this 40% failure rate is typical of best clinical practice (7). It would appear that the reduction in endothelial injury (i.e shear stress) and enhanced flow stasis (i.e. infusion dynamics) due to the ramped tip design contributed to the improved outcomes.

Recent clinical trial data shows that device failure remains unacceptable despite best efforts by end-users. In the SAVE Trial (23), approximately 40% of PIVCs failed in each arm of the study. In a trial investigating post-insertion flushing (24), failure was 22% in the intervention group compared to 30% in controls. Most recently, in the PREBACP trial (7), 37% of devices in the intervention group failed; an improvement of about 9%. Therefore, at best, one in five PIVCs fail, and at worst, two in five. Improvements in the PIVC design are likely to help reduce these rates, and the FloRamp may be a step towards this goal.

We designed an experimental protocol that mimicked current best clinical practice, including the use of a continuous infusion and regular PIVC flushes, and aimed to minimise any confounding factors on failure so that only the impact of the PIVC design was under evaluation. We ensured the device could not be dislodged by sedating the animals and used a contemporary ‘care bundle’ of aseptic touch, sterile dressings, securement of device, and frequent monitoring and measurement of VIP score. For each flush procedure, we measured the resistance, and compared to the control device, we found that the FloRamp enabled resistance-free infusion for longer before any noticeable resistance occurred. We observed only two FloRamp devices (2/5) to register a VIP score, compared to four control devices (4/5); however, the two FloRamp devices did begin to show signs of early phlebitis at 30 h and 48 h. This reduction in VIP has important implications as the VIP is a simple marker used widely in clinical practice. Five of the seven histopathology markers we assessed were lower in veins inserted with the FloRamp, although no comparisons were statistically significant. Of particular note is the exfoliation of endothelium and intimal proliferation, oedema and necrosis; the FloRamp was purposely designed to reduce endothelial shear stress acting on the intima of the vein, with levels above 38 Pa known to strip endothelial cells from the intimal layer (25). Furthermore, the effects of shear stress on platelet adhesion and endothelial activation (26-28) are well known and contribute to inflammation, coagulation and potentially PIVC failure. By reducing the shear stress, the FloRamp reduces the critical instigator of phlebitis (assessed here by VIP score and mural inflammation), which in turn down-regulates the processes leading to intimal injury and vessel occlusion.

We followed the histopathological methods reported in a previous study investigating a novel PIVC design. Weiss et al. (19) also inserted 20 G BD Insyte PIVCs into the veins of porcine ears, however their study lasted 12 days, compared to our 2-day study, with typical clinical practice indicating PIVC usage for short-term administration of fluids and medication (2-3 days) in critical conditions requiring mechanical ventilation and critical care management. After 12 days from cannulation, Weiss et al. found much greater intimal proliferation, intimal oedema, and mural haemorrhage, whereas the control devices in our study exhibited much greater mural inflammation, exfoliation of endothelium and intimal necrosis. These comparisons to our study are interesting; our sedated pigs were studied for a much shorter time, whereas the pigs in their study were free to move about, eat and drink, with the authors noting that the pigs were “constantly trying to remove the devices.” The authors also state that the securement devices used did not eliminate relative motion between the catheter and vein. Nevertheless, Weiss et al. report much lower inflammation and general tissue damage. It is important to emphasise that in our study we applied inclusive qualitative assessment of injury, i.e. VIP, which resulted in injury dynamics in line with clinical observations in humans.

A key advantage of the FloRamp is its simplicity as it is a relatively straightforward dimension change that captures important engineering and biomedical principles; yet heralds a significant change in the PIVC design, ameliorating some of the impact of cannulation and infusion. This subtle design change lends itself to the regulatory process involved in medical devices, as well as the scale at which PIVCs are produced. Furthermore, as it is ergonomically identical to a typical PIVC, there is no additional end-user training required to use in practice.

Although there are many strengths to our study, there are also some notable limitations. First, this was a preliminary study of only five vascular access sites in four pigs, and the small sample size should be considered while interpreting performance of the novel device, in particular, the statistical comparisons between devices. Second, as two control devices occluded halfway through the experiment, we excluded those veins from the quantitative histological analysis. This further reduced the samples for histological comparison. Third, our 48 hour experiment was relatively short and in a future study the duration should be extended until at least 72 hours, or preferably, only terminated on failure. Fourth, while the pig is a useful model for vascular access, there are differences in anatomy, with the fatty subdermal layer of the skin being thicker in pigs, and therefore studying the FloRamp in humans is the next logical step. Finally, this was a well-controlled animal study using sedated pigs and while the protocol may resemble some clinical scenarios such as intensive care, the findings may not translate to general PIVC usage.

In conclusion, despite best efforts from end users, premature failure of PIVCs remains a significant healthcare problem. Here we report a new PIVC design that appears to hold promise, outperforming the current state-of-the-art in a well-controlled study using an animal model of vascular access. The FloRamp specifically addresses calls from the vascular access community for PIVC producers to revisit PIVC design and may be a useful new design feature for PIVC manufacturers to incorporate into devices.

## ACKNOWLEDGEMENTS

The authors acknowledge the efforts of Dr Roland Steck and the staff of Queensland University of Technology’s Medical Engineering Research Facility for their support and assistance for this study. The authors also acknowledge Dr Felicity Lawrence and Dr John Mackie from QML Vetnostics for their assistance with the histopathology. The authors would also like to thank A/Prof Andrew Bulmer from Griffith University for his valuable comments and insights into the work.

## AUTHOR CONTRIBUTIONS

BJD and CS conceived the study. LJK and BJD performed computer simulations. BJD, CS, GLB and SK designed the animal study. GA, CA, NS, SL, MB, MRP, ESW, KS, KL, SH, KW and GLB performed the animal experiments. BJD interpreted the data, discussed the results with all authors, and wrote the manuscript, which was then reviewed and edited by all authors. All authors approved the final manuscript for submission.

## REFERENCES

1. Helm RE, Klausner JD, Klemperer JD, Flint LM, Huang E. Accepted but unacceptable: peripheral IV catheter failure. J Infus Nurs. 2015;38(3):189–203.

2. Chen S, O’Malley M, Chopra V. How common are indwelling devices in hospitalized adults? A contemporary point prevalence study in a tertiary care hospital. American Journal of Infection Control. 2021;49(2):194–7.

3. Burnaby B. Global market overview for vascular acceess devices and accessories 2012–2022. 2017. Contract No.: 4339819.

4. Verified Market Research. Global Peripheral IV Catheter Market: Market Size, Status and Forecast to 2028. 2020.

5. Marsh N, Webster J, Larson E, Cooke M, Mihala G, Rickard CM. Observational Study of Peripheral Intravenous Catheter Outcomes in Adult Hospitalized Patients: A Multivariable Analysis of Peripheral Intravenous Catheter Failure. J Hosp Med. 2018;13(2):83–9.

6. Ray-Barruel G, Xu H, Marsh N, Cooke M, Rickard CM. Effectiveness of insertion and maintenance bundles in preventing peripheral intravenous catheter-related complications and bloodstream infection in hospital patients: A systematic review. Infection, Disease & Health. 2019;24(3):152–68.

7. Blanco-Mavillard I, de Pedro-Gómez JE, Rodríguez-Calero M, Bennasar-Veny M, Parra-García G, Fernández-Fernández I, et al. Multimodal intervention for preventing peripheral intravenous catheter failure in adults (PREBACP): a multicentre, cluster-randomised, controlled trial. Lancet Haematol. 2021;8(9):e637–e47.

8. Zdrahala RJ, Spielvogel DE, Strand MA. Softening of Thermoplastic Polyurethanes: A Structure/Property Study. Journal of Biomaterials Applications. 1987;2(4):544–61.

9. Kuş B, Büyükyılmaz F. Effectiveness of vialon biomaterial versus teflon catheters for peripheral intravenous placement: A randomized clinical trial. Jpn J Nurs Sci. 2020;17(3):e12328.

10. González López JL, Arribi Vilela A, Fernández del Palacio E, Olivares Corral J, Benedicto Martí C, Herrera Portal P. Indwell times, complications and costs of open vs closed safety peripheral intravenous catheters: a randomized study. J Hosp Infect. 2014;86(2):117–26.

11. Kumar DR, Hanlin E, Glurich I, Mazza JJ, Yale SH. Virchow’s Contribution to the Understanding of Thrombosis and Cellular Biology. Clinical Medicine & Research. 2010;8(3-4):168–72.

12. Hawthorn A, Bulmer AC, Mosawy S, Keogh S. Implications for maintaining vascular access device patency and performance: Application of science to practice. The Journal of Vascular Access. 2019;20(5):461–70.

13. Traub O, Berk BC. Laminar Shear Stress. Arteriosclerosis, Thrombosis, and Vascular Biology. 1998;18(5):677–85.

14. Hathcock JJ. Flow effects on coagulation and thrombosis. Arterioscler Thromb Vasc Biol. 2006;26(8):1729–37.

15. Piper R, Carr PJ, Kelsey LJ, Bulmer AC, Keogh S, Doyle BJ. The mechanistic causes of peripheral intravenous catheter failure based on a parametric computational study. Scientific Reports. 2018;8(1):3441.

16. Doyle B, Kelsey L, Carr PJ, Bulmer A, Keogh S. Determining an Appropriate To-Keep-Vein-Open (TKVO) Infusion Rate for Peripheral Intravenous Catheter Usage. Journal of the Association for Vascular Access. 2021;26(2):13–20.

17. Chistiakov DA, Orekhov AN, Bobryshev YV. Effects of shear stress on endothelial cells: go with the flow. Acta Physiol (Oxf). 2017;219(2):382–408.

18. Eisa-Beygi S, Benslimane FM, El-Rass S, Prabhudesai S, Abdelrasoul MKA, Simpson PM, et al. Characterization of Endothelial Cilia Distribution During Cerebral-Vascular Development in Zebrafish (Danio rerio). Arterioscler Thromb Vasc Biol. 2018;38(12):2806–18.

19. Weiss D, Yaakobovitch H, Tal S, Nyska A, Rotman OM. Novel short peripheral catheter design for prevention of thrombophlebitis. J Thromb Haemost. 2019;17(1):39–51.

20. Jackson A. Infection control--a battle in vein: infusion phlebitis. Nurs Times. 1998;94(4):68,71.

21. Shackelford C, Long G, Wolf J, Okerberg C, Herbert R. Qualitative and Quantitative Analysis of Nonneoplastic Lesions in Toxicology Studies. Toxicologic Pathology. 2002;30(1):93–6.

22. Marsh N, Rickard CM. Peripheral intravenous catheter failure-is it us or is it them? Lancet Haematol. 2021;8(9):e615–e7.

23. Rickard CM, Marsh N, Webster J, Runnegar N, Larsen E, McGrail MR, et al. Dressings and securements for the prevention of peripheral intravenous catheter failure in adults (SAVE): a pragmatic, randomised controlled, superiority trial. The Lancet. 2018;392(10145):419–30.

24. Keogh S, Shelverton C, Flynn J, Mihala G, Mathew S, Davies KM, et al. Implementation and evaluation of short peripheral intravenous catheter flushing guidelines: a stepped wedge cluster randomised trial. BMC Medicine. 2020;18(1):252.

25. Fry DL. Acute vascular endothelial changes associated with increased blood velocity gradients. Circ Res. 1968;22(2):165–97.

26. Dunkley S, Harrison P. Platelet activation can occur by shear stress alone in the PFA-100 platelet analyser. Platelets. 2005;16(2):81–4.

27. Ploppa A, Schmidt V, Hientz A, Reutershan J, Haeberle HA, Nohé B. Mechanisms of leukocyte distribution during sepsis: an experimental study on the interdependence of cell activation, shear stress and endothelial injury. Crit Care. 2010;14(6):R201.

28. Galbusera M, Zoja C, Donadelli R, Paris S, Morigi M, Benigni A, et al. Fluid shear stress modulates von Willebrand factor release from human vascular endothelium. Blood. 1997;90(4):1558–64.

